# Cell-type heterogeneity in adipose tissue is associated with complex traits and reveals disease-relevant cell-specific eQTLs

**DOI:** 10.1101/283929

**Authors:** Craig A. Glastonbury, Alexessander Couto Alves, Julia S. El-Sayed Moustafa, Kerrin S. Small

## Abstract

Adipose tissue is comprised of a heterogeneous collection of cell-types which can differentially impact disease phenotypes. We investigated cell-type heterogeneity in two population-level subcutaneous adipose tissue RNAseq datasets (TwinsUK, N =766 and GTEx, N=326). We find that adipose cell-type composition is heritable and confirm the positive association between macrophage proportion and obesity (BMI), but find a stronger BMI-independent association with DXA-derived body-fat distribution traits. Cellular heterogeneity can confound ‘omic analyses, but is rarely taken into account in analysis of solid-tissue transcriptomes. We benchmark the impact of adipose tissue cell-composition on a range of standard analyses, including phenotype-gene expression association, co-expression networks and *cis*-eQTL discovery. We applied G x Cell Type Proportion interaction models to identify 26 cell-type specific eQTLs in 20 genes, including 4 autoimmune disease GWAS *loci*, demonstrating the potential of *in silico* deconvolution of bulk tissue to identify cell-type restricted regulatory variants.

## Introduction

Adipose tissue is the largest endocrine organ in the human body and has a role in the development of insulin resistance, cardiovascular disease, type 2 diabetes and many other cardiometabolic disorders. Adipose tissue is heterogeneous, it is comprised of an array of cell types including adipocytes, pre-adipocytes, endothelial cells, and several immune cell subtypes (Rafols 2014). Adipose tissue cellular composition changes in response to obesity and it is thought that this change, in particular the marked increase in immune cell infiltration, contributes to some of the negative health consequences of obesity (Chawla et al., 2011; Cancello et al., 2005; Heilbronn et al., 2008). It is therefore of interest to understand the cellularity of adipose tissue, its variability in the population and how this affects health and disease.

Due to the biomedical importance and relatively easy physical accessibility of subcutaneous adipose tissue, a large body of adipose transcriptomic datasets has been generated from healthy volunteers and patients, including several studies with greater than 200 participants (Grundberg *et al*., 2012; Buil et al., 2015; Emilsson et al., 2008; Greenawalt et al., 2011; Civelek et al., 2017; Lonsdale et al., 2013). To our knowledge, these studies have not assessed the cellular composition of their samples, despite the fact that cellular heterogeneity is a well-established confounder in transcriptomic analysis of bulk tissues (Titus et al., 2017; Lappalainen et al., 2017; Jaffe et al., 2014). While extensive investigation and methodological development has centered on computationally accounting for cell-type composition in whole blood (Jaffe et al., 2014), very few studies have investigated the extent of cellular heterogeneity in other tissues and how it impacts –omic level analyses (McCall et al., 2016). Directly assessing cell composition in adipose is challenging, methods such as flow sorting face technical difficulties including adipocyte rupturing and shared cell-type specific surface markers, as well as low-throughput and high expense when applied to hundreds of samples. Single-cell analysis could overcome some of these considerations, however the complex logistics of population level collection of adipose biopsies and expense of single cell analysis mean that there is considerable utility of *in silico* deconvolution of ‘omic profiles generated from bulk adipose tissue.

Here, we utilize *in silico* deconvolution to estimate the relative proportions of four distinct cell types (adipocytes, macrophages, CD4+ t-cells and Micro-Vascular Endothelial Cells (MVEC)) in bulk subcutaneous adipose tissue transcriptomes from two independent datasets; 766 individuals from TwinsUK and 326 post-mortem GTEx donors. We conduct extensive simulations to investigate whether our methods accurately identify the relevant cell types, the range of cell type detection possible and robustness to varying levels of noise and unknown cell content (contamination). We find significant cellular heterogeneity within and between these datasets. We recapitulate the well-known cellular hallmark of obesity, finding a positive association between adipose tissue macrophage abundance and Body Mass Index (BMI), but identify stronger relationships to DXA-derived body fat distribution traits. We assess the impact of adipose cellular heterogeneity on standard ‘omic analyses, including *cis*-eQTL discovery, co-expression networks and differential gene expression studies. Finally, we utilize cell-type composition in interaction models to identify cell-type specific eQTLs from bulk tissue transcriptomes that are enriched for GWAS variants and cell type relevant enhancers.

## Results

### Accurate cell type estimates that are robust against unknown content and noise

We estimated cell-type proportion in bulk adipose tissue RNA-seq profiles with CIBERSORT, a ν-support vector regression (ν-SVR) method developed to estimate cell proportions using gene expression obtained from solid tissues (Newman et al. 2015). CIBERSORT identifies cell-type specific marker genes from purified cell transcriptome profiles to construct a tissue-specific signature matrix, a set of differentially expressed genes across all cell types and utilizes this signature matrix to perform the deconvolution step.

To construct the CIBERSORT adipose tissue signature matrix, we obtained previously published RNA-seq datasets from purified cells that are known to be present in subcutaneous adipose tissue including Adipocytes, Macrophages, CD4+ T-cells and microvascular endothelial cells (MVEC) (**Table: S1**). Adipose tissue is comprised of many more cell types than the four we focus on here. Estimation of additional cell types was prevented by either the unavailability of purified RNA-seq datasets for those cell types at the time of study, lack of replicates to ensure stable construction of the signature matrix or very low frequency in the tissue (such as mast cells). Biological replicates of each of the four purified cell type were included. Hierarchical clustering of the purified cell transcriptional profiles recapitulated developmental cellular hierarchy (**Figure S1**). The final CIBERSORT adipose signature matrix is comprised of 658 genes including several well-known cell-type specific markers such as *SCD, COL1A1* and *ADIPOQ* in adipocytes, *SERPINE1, MMP1* and *VWF* in endothelial cells, *SPP1, F13A1* and *CTSC* in Macrophages and *FOS, TCF7* and *CD3* in t-cells. The full signature matrix is included in **Supplementary File 1**.

To test deconvolution ability, accuracy and robustness to noise, we performed several typical simulations used to benchmark deconvolution accuracy (Newman et al., 2015; Gong & Szustakowski 2013). First, we tested whether the adipose tissue signature matrix can accurately identify the four cell types when applied to a set of independent, purified cell type RNA-seq datasets). All benchmark cell types were estimated with high accuracy, with three out of four cell types attaining ≥ 99% accuracy in prediction (**Table S2**. Macrophages (93% accuracy) are particularly difficult to purify, so it is possible that the 6% CD4+ t-cells we estimate in the macrophage benchmark sample were present in the original purified macrophage sample.

We next assessed ability to estimate the constituent cell proportions of a mixture of known cell types. We created 1000 *in-silico* mixtures of known proportions of each of the four cell types with the independent purified cell type datasets (**Table S1**), Application of CIBERSORT to the *in silico* mixtures yielded highly accurate estimates of cell type proportion, with mean absolute deviation (mAD) of estimated proportions to ground truth values ranging from 0.019 to 0.068 (**Figure 1**). Biopsies can contain contaminant cells from other tissues, which could impact cell-type proportion estimates if contaminant cells share marker genes with any of the four cell types we are estimating. To test this, we added proportions of smooth muscle cells, dendritic cells and neutrophils to the *in-silico* mixtures of the four cell types. These cell types can be present in adipose tissue and therefore reflect realistic ‘contaminant cells’. Neutrophils in particular could be troublesome, as neutrophils make up 60-70% of whole blood, and whole blood may be present in the biopsy at varying levels as a contaminant. Cell type prediction was accurate when up to 10% of a sample was composed of contaminant cell types (**Figure S2**). As it is likely the content of unknown cells in the samples is ≤ 5% given previously published cell type estimates from adipose tissue (Travers et al., 2015; Zimmerlin et al., 2010; Van Harmelen et al., 2003), the adipose tissue signature matrix is robust in estimating cell types from mixtures with some unknown content.

**Figure 1:**
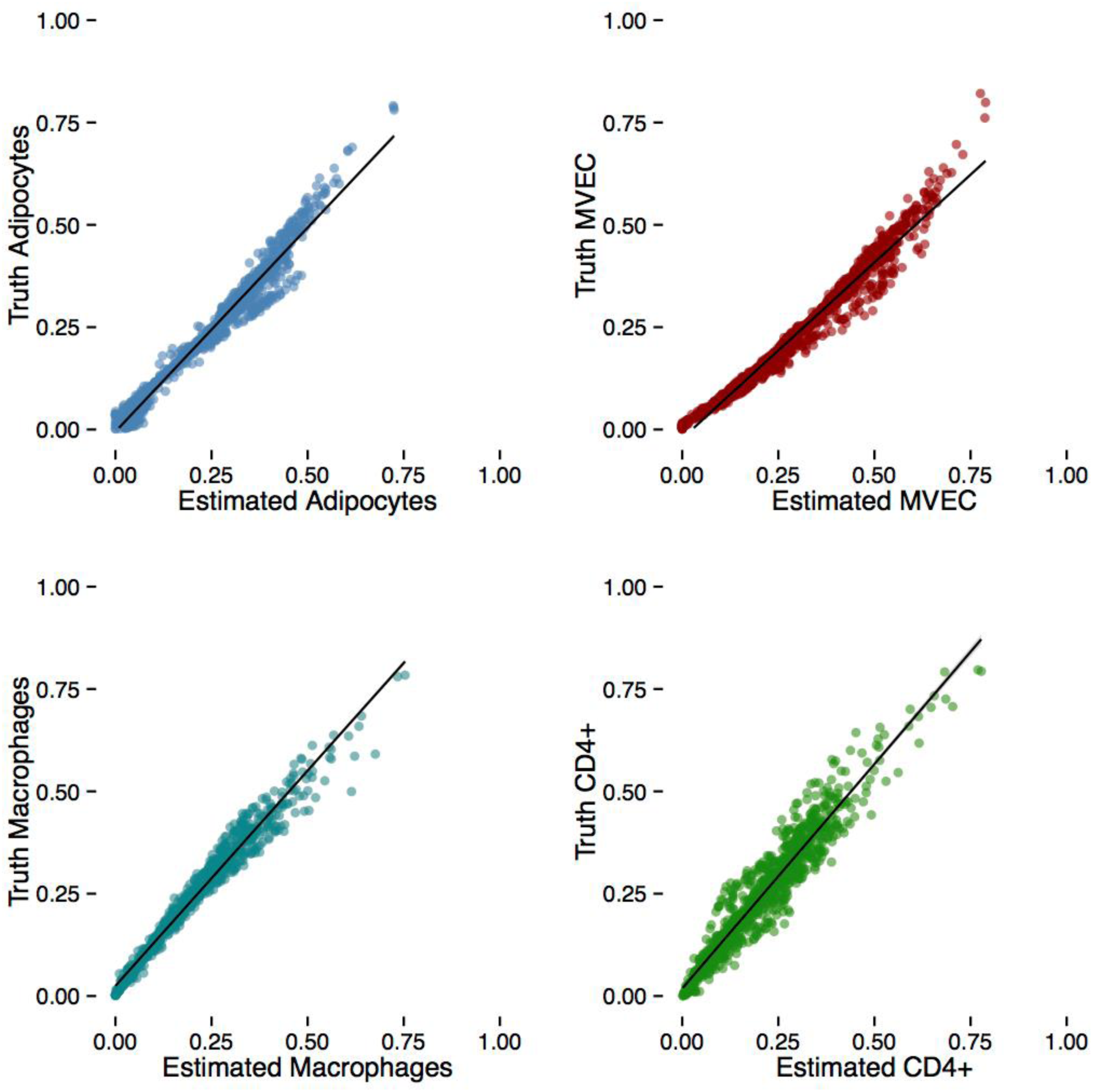
Cell-type proportion is accurately estimated in *in silico* mixture simulations. Each panel displays cell-type estimation in 1000 in silico mixtures. Each point represents one simulation.

Finally, given technical factors during library preparation and sequencing introduce noise in RNA-seq experiments, we tested how much noise we could introduce into the simulations and still accurately predict cell type proportions, similar to analyses performed in Newman et al. (2015). We added Gaussian noise at 10, 50 and 90% as well as the naturally occurring noise present in each of the separate experimentally derived purified RNA-seq datasets. The estimates are robust when up to 10% of the mixture is distorted with noise, and a linear relationship between ground truth and predicted estimates still holds when large amounts of noise are introduced (**Figure S3**).

In summary, we have adapted CIBERSORT to accurately predict cell-type composition in bulk RNA-seq data from adipose tissue. Under reasonable limits of contamination and noise our estimates of adipose tissue cell-types proportion are reliable.

### Estimation of relative cell type proportions in bulk adipose RNAseq datasets

We applied CIBERSORT and the adipose tissue signature matrix to a previously published dataset of 766 subcutaneous adipose tissue biopsies obtained from female twin participants of TwinsUK (Grundberg et al., 2012 Buil et al., 2015) All 766 TwinsUK RNA-seq samples were successfully deconvolved at an FDR 1%. Adipocytes were the most dominant relative cell-type [0.73-0.99], but also show significant inter-subject variability (**Figure 3**). Proportions of the other estimated cell types ranged from [0.004-0.22] for macrophages (M1/M2 combined), [0-0.19] for MVEC and [0-0.11] for CD4+ t-cells (**Figure 3**). These estimates agree with previously published studies performed using flow cytometry (**Supplemental Table S3**). As the vast majority of TwinsUK adipose samples had CD4+ t-cell estimates below 1%, we chose to focus on adipocytes, macrophage and endothelial cell estimates for downstream analysis.

**Figure 2:**
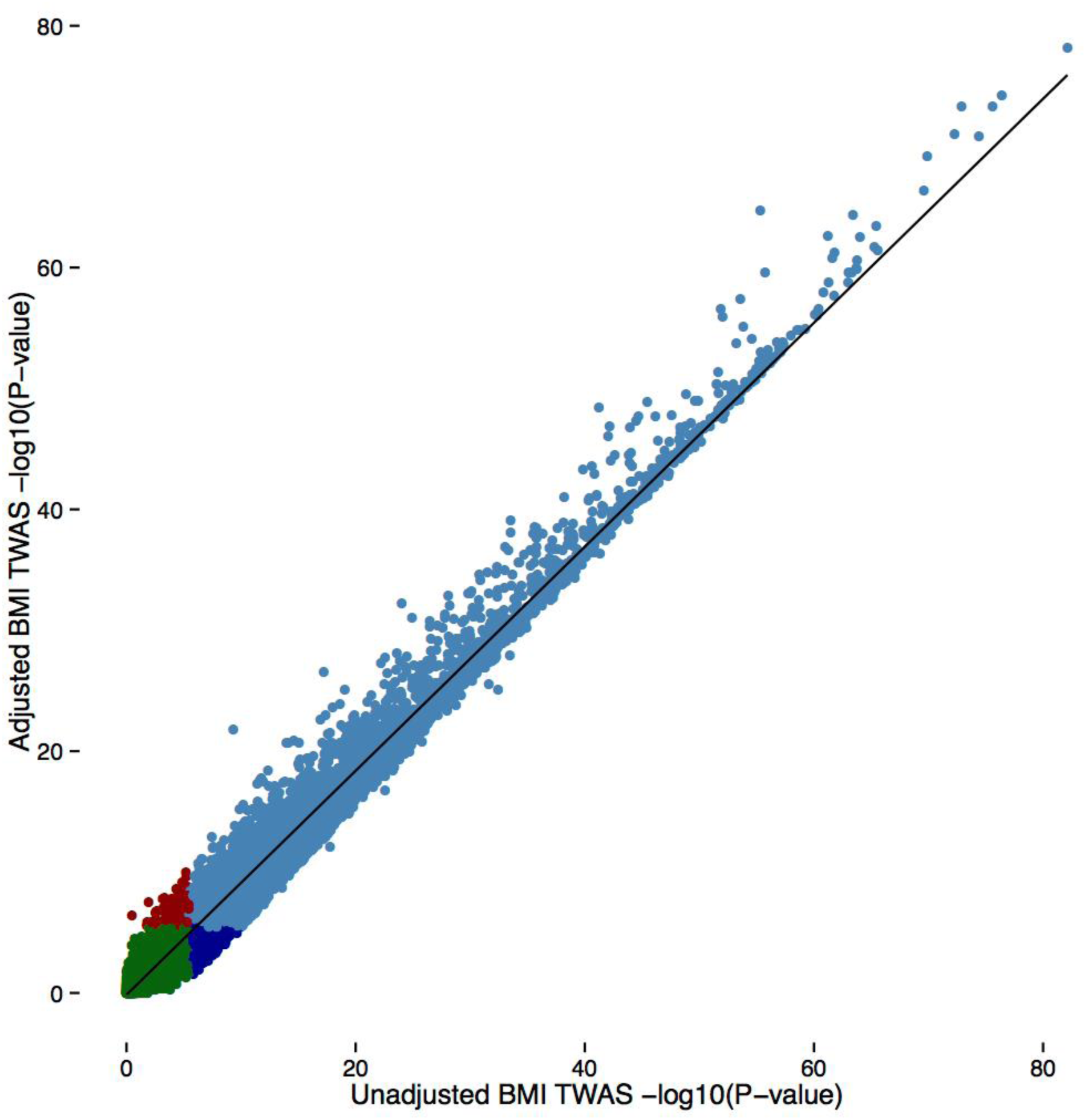
Adjusting for macrophage proportion accounts for 11% of gene-BMI associations. Each point represents one gene and is coloured as follows: Red – significant in macrophage adjusted TWAS only, dark blue significant in unadjusted TWAS only, light blue – significant in both TWAS and green - not-significant in both TWAS.

**Figure 3:**
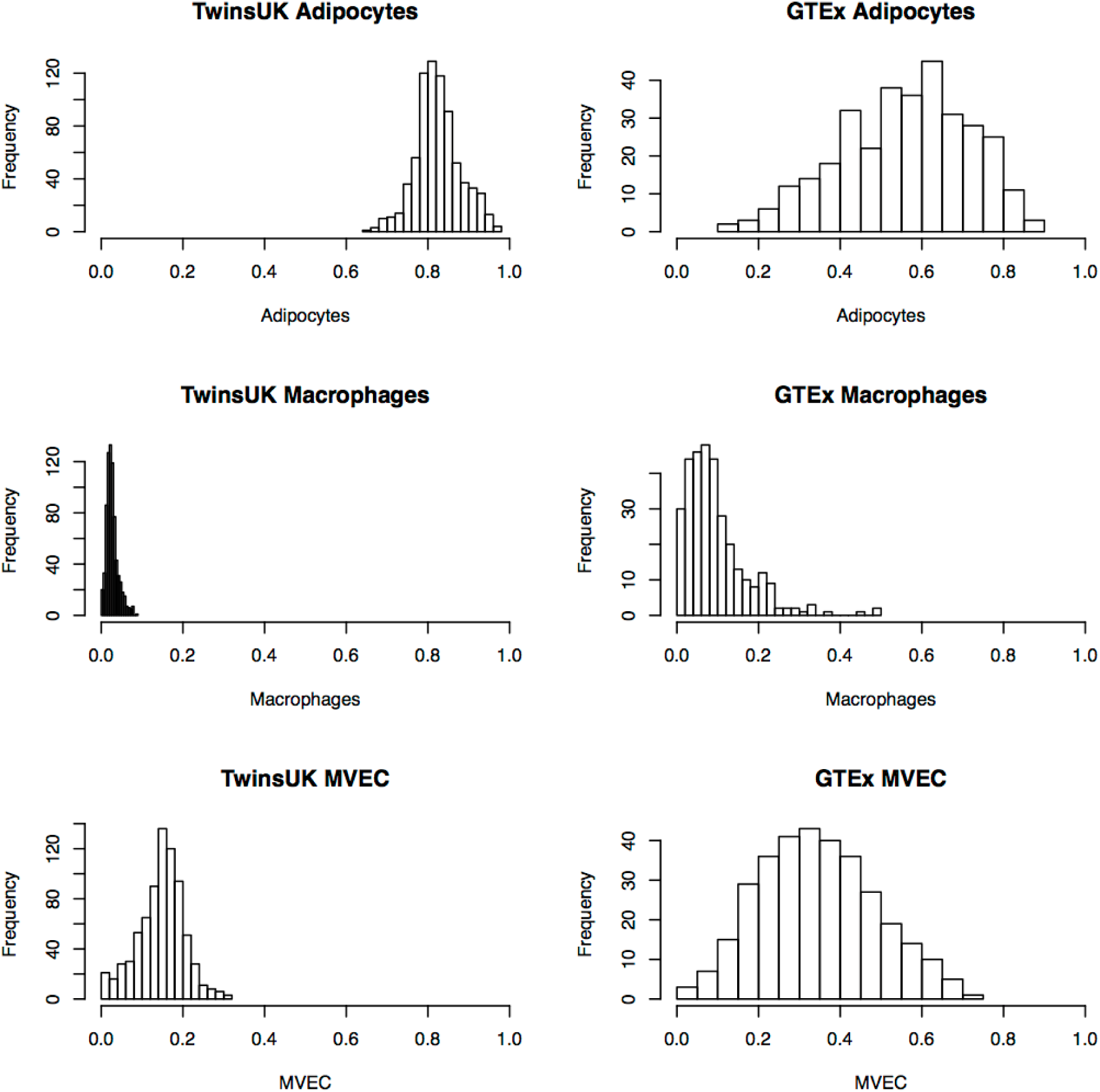
Distribution of relative cell type estimates in TwinsUK and GTEx adipose samples. TwinsUK samples are shown on the left, GTEx on right.

We next applied CIBERSORT to an independent sample of 326 post-mortem subcutaneous adipose tissue biopsies from the genotype tissue-expression consortium (GTEx) (Aguet et al., 2017). In contrast to TwinsUK, ∼23% of GTEx samples (75/326) failed successful deconvolution (1% FDR), suggesting substantial differences of cell types present in the tissue from the signature matrix. The 251 GTEx samples that passed deconvolution had markedly different cell-type composition profiles as compared to the TwinsUK samples, with lower adipocyte fraction (GTEx_median_ = 0.62, TwinsUK_median_ = 0.82), twice as much vasculature (GTEx median MVEC proportion = 0.30, TwinsUK = 0.15) and 4 times as many macrophages (GTEx median macrophage = 0.08, TwinsUK = 0.02) (**Figure 3**). To assess the GTEx estimates, we investigated whether there were visible histological differences between samples with predicted extreme macrophage proportion estimates in histology slides of the GTEx biopsies. We observed concordance between our estimates and visual inspection of the histology slides. We demonstrate this in **Figure 4**, where the sample with the lowest macrophage proportion (0%) is composed of adipocytes with few additional cells present and the sample with the highest macrophage proportion (49%) has a substantial amount of vasculature/blood cells surrounding the adipocytes.

**Figure 4:**
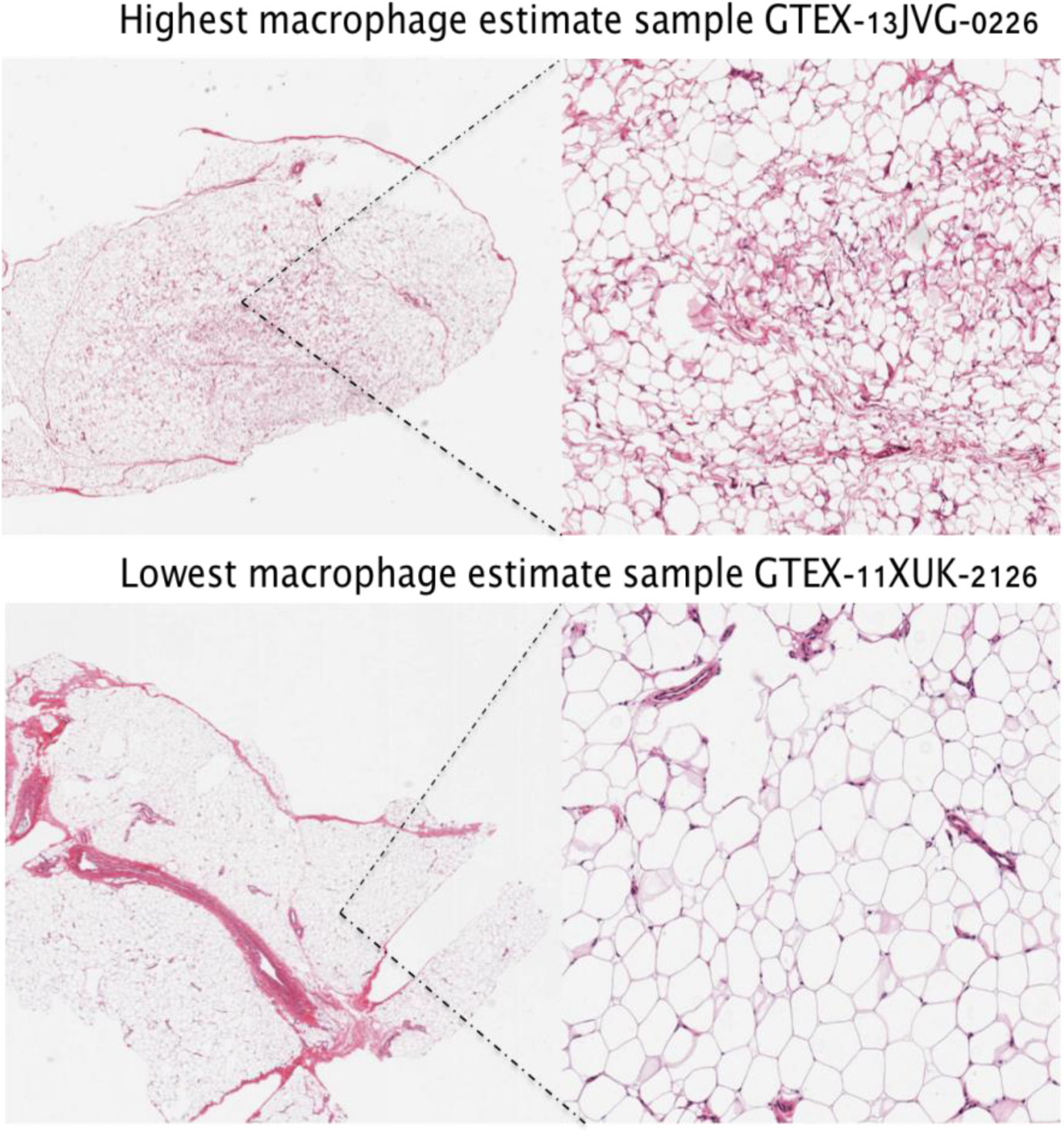
Estimated cell-type composition of GTEx samples corresponds to image data. Histology images from the GTEx adipose samples with highest (49%) (top) and lowest (0%) (bottom) macrophage estimates are shown. Both whole biopsy (left) and zoomed in images (right) are presented. Estimated cell type composition of all GTEx samples are provided in Supplementary File 2.

To validate the difference in adipocyte proportion between the datasets, we focused on the expression of *ADIPOQ*, which encodes the hormone adiponectin. *ADIPOQ* expression is expressed highly in adipocytes and pre-adipocytes. *ADIPOQ* was expressed 4-fold higher in TwinsUK (median TMM = 3998, expression rank = 19) compared to GTEx (median TMM = 963, expression rank = 59). The distribution and range of *ADIPOQ* expression varied between the datasets, following a normal distribution in TwinsUK (untransformed, TMM data) and Poisson in GTEx (**Supplementary Figure S8**). *ADIPOQ* expression is very low in some GTEx samples as compared to TwinsUK, which suggests fewer viable adipocytes (GTEx *ADIPOQ*_*min*_ = 3.13 TMM; TwinsUK *ADIPOQ*_*min*_ = 986 TMM) (**Figure S6**). The *ADIPOQ* results strongly support the CIBERSORT estimates of lower adipocyte proportion in the GTEx samples.

There are several possible explanations of why cell-type proportions differ between TwinsUK and GTEx. The function and metabolic activity of adipose tissue is known to vary between fat depots – markedly between android and gynoid depots. The gynoid GTEx adipose samples were obtained via surgical incision from the lower left leg, whilst android TwinsUK samples were derived from punch biopsies from the abdomen. Fibrosis due to ischemia is likely to alter the number of viable cells available for sequencing in post-mortem samples; GTEx pathologist notes frequently recorded the presence of large fibrotic regions (up to 60% of a given histology slide) in addition to contaminant cells such as nerve tissue, bone ossification or the presence of large blood vessels. Given the large disparities in cell composition between the datasets we chose to focus on the TwinsUK dataset for the following analysis, both for the availability of additional phenotypes and as their composition reflects *in vivo* ranges.

### Adipose cell-type proportions are heritable

Several studies have demonstrated that the cell-type composition of whole blood is heritable (Roederer et al., 2016, Brodin et al. 2015), but the influence of genetics on adipose cell-type composition has not been explored. Using classical twin models, we estimate the narrow sense heritability (*h*^*2*^) of adipocyte, macrophage and endothelial cell proportion to be 17%, 30% and 21% respectively in the TwinsUK data. The heritability of adipose tissue cell composition may be mediated by genetic drivers of whole-body traits such as BMI that in turn drive changes in cell-type proportion, or may be mediated by local effects within adipose tissue such as rates of adipo- or angiogenesis.

### Adipose tissue cell type proportion is associated to whole-body obesity traits but not age

Macrophage infiltration and increased abundance in adipose tissue is known to increase with obesity and its associated chronic inflammation (Boutens et al., 2016). We recapitulate this finding, demonstrating a significant correlation between Body Mass Index (BMI) and estimated adipose macrophage proportion in TwinsUK (**Table: 1**). We explored the relationship between cell-type composition and body-fat distribution using highly accurate Dual X-ray absorptiometry (DXA) measures of visceral fat volume (VFAT) and android/gynoid (A/G) ratio in a subset of twins (N = 652) with concurrently measured DXA scans. Despite the smaller sample size, the correlation of android/gynoid ratio and visceral fat with relative macrophage estimates were significantly higher than to BMI (**Table: 1**). Including BMI as a covariate did not change the associations to DXA-derived traits, indicating body-fat distribution is associated to adipose tissue cell composition independent of overall adiposity. This finding confirms both the importance of macrophage biology in obesity but also suggests inflammation plays a more prominent role in body-fat distribution than currently appreciated.

In contrast to the well-documented association between whole blood cell-type composition and age (Jaffe & Irizarry 2014), there was no association between age and either macrophage or adipocyte proportion (r = −0.02). This indicates that adipose cell-type composition is not a major confounder in identification of age-related transcripts (Viñuela et al., 2017) or differentially methylated regions (Nilsson et al., 2014) in adipose tissue.

### Adjusting for Macrophage heterogeneity accounts for 11% of BMI differentially expressed genes

BMI has a profound effect on adipose tissue gene expression; the majority of the adipose transcriptome is associated to BMI in studies conducted with both microarrays and RNA-seq in independent populations (Emilsson et al. 2008, Glastonbury *et al*. 2016, Civelek et al., 2017). It is unclear how much of BMI-associated changes in gene expression are mediated by the changes in cell-type composition that accompany increasing BMI. To address this, we identified associations between gene expression and BMI under two models, adjusting and not adjusting for macrophage proportion. In the first model, we recapitulate previous results with expression of 6,366/14,897 protein-coding genes significantly associated with BMI (Bonferroni corrected *P-value* = 3.56 × 10^-6^). Adjusting for macrophage proportion resulted in 11% of associations no longer being significant, a loss of 707 gene expression-BMI associations (**Figure 2**). This demonstrates that whilst inflammation is an important aspect of obesity etiology, the majority (89%) of BMI-expression associations are likely to be independent of macrophage proportion, but could still be dependent on activation state. Of the 707 genes that are no longer significant after adjusting for macrophage proportion an example is *CD209* P-value_original_ = 7.72 × 10-8, P-value_adj_ = 0.0019), a gene that encodes for a C-type lectin that is found primarily on the surface of macrophages and dendritic cells. Additional example genes no longer significant include *LILRA2, MNDA* and *CMKLR1* that are known to be primarily expressed in macrophage and immune cell lineages (Lee et al., 2007; Briggs et al., 1994; Zabel et al., 2006).

### Cell-type proportion explains major components of gene expression variance and co-variance

Principle component analysis (PCA) is commonly used to understand the sources of gene expression variance. We identified principle components in the TwinsUK samples and correlated them to cell-type proportion estimates. PC1 was correlated with adipocyte and endothelial cell proportion (R = 0.40, P-value ≤ 2.2 × 10^-16^; R = 0.41, P-value = 2.2 × 10^-16^, respectively). PC2 was negatively correlated with macrophage proportion (R = −0.63, P-value ≤ 2.2×10^-16^ and positively correlated with endothelial cell proportion (R = 0.21, P-value ≤ 3.7 × 10^-9^). PC1 and PC2 cumulatively explained 25% of adipose tissue gene expression variance (**Figure S6**). This indicates that cell-type heterogeneity at the population level is a major driver of gene expression variation in adipose tissue, and accounting for principle components in downstream analysis should account for some of this variability.

Weighted Gene Co-Expression Network Analysis (WGCNA) is a widely used technique that uses the correlation structure of global gene expression profiles to construct modules of genes, some of which have been ascribed distinct functional roles or correspond to gene networks. 11 out of 13 WCGNA modules in TwinsUK correlated with cell-type proportion (**Figure S7**, *P-value* < 0.0038). The most significant macrophage-proportion associated module (Pearson’s R = 0.67, *P-value* ≤ 2.2× 10^-16^) (**Figure S7a**), recapitulated the Macrophage Enriched Metabolic Network (MEMN), an adipose gene expression signature associated with increasing BMI (Emilsson et al. 2008, Chen et al. 2008). The MEMN-green module’s constituent genes were enriched for glycoproteins (*P-value* = 7.1 × 10^-63^), Immunity (*P-value* = 1.1 × 10^-23^) and the innate immune response (*P-value* = 4.5 × 10^-12^). Endothelial cell proportion was positively correlated to the turquoise module (R = 0.41) which was significantly enriched for GO terms related to angiogenesis (*P-value* = 6.4 × 10^-12^). These findings demonstrate that cell-type composition is a major driver of co-expression in bulk tissue RNA-seq samples and could confound analysis if samples are not matched for cell-type proportion.

### Correction for macrophage heterogeneity in adipose tissue increases *cis*-eQTL discovery yield

To determine if adipose cell-type heterogeneity can confound *cis*-eQTL analysis we investigated the effect of correcting for cell-type in *cis*-eQTL analysis. We implemented a naive *cis*-eQTL discovery model, not adjusting for any cell-type proportion and a separate, macrophage-corrected eQTL model. Adjusting for macrophage heterogeneity amongst samples led to a modest increase in *cis*-eQTL yield (2.3%) (naive model = 5,531, macrophage-adjusted = 5,665 SNP-gene pairs, FDR5%). However, it has become standard practice in *cis*-eQTL studies to use gene expression principle components, PEER factors, or other factor-analysis based methods, to estimate and adjust-out confounding factors from gene expression data. To test whether latent factors account for cell-type proportion variability, we re-ran the naive and cell-type adjusted *cis*-eQTL scans including adjustment for 30 PEER factors. PEER factor adjustment achieved a similar increase in *cis*-eQTL yield in the naïve and cell-adjusted models respectively and resulted in near identical results (naïve-PEER = 7,665, macrophage-adjusted-PEER = 7,664). This confirms that latent factors capture the cell type composition differences amongst adipose samples, as well as many other unmeasured latent factors, but if covariates are known, it is better to adjust using a full specified model rather than estimate latent factors given the known risk of collider bias (Dahl et al., 2017).

### Identification of cell-type specific eQTLs from bulk tissue

Previous studies have identified cell-type specific eQTLs in bulk whole blood expression profiles by fitting G x cell type proportion interaction models (Westra et al. 2015). We utilized this strategy to detect cell-type specific *cis*-eQTLs in the TwinsUK adipose data. At a strict Bonferroni-corrected threshold (*P-value* threshold = 1.01 × 10^-9^, based on 49,219,795 association tests in the 1 MB TSS centered window around 14,897 genes) we identified 26 G x cell type interactions at 20 unique genes (**Table 2**). Twelve gene-SNP pairs had an interaction with macrophage proportion, 10 with endothelial proportion and four with adipocyte proportion (**Table 2**). Examples include *MARCO*, a macrophage receptor with collagenous structure, whose expression depends on macrophage proportion and rs1884841. *TC2N*, which is responsible for the secretion of *VWF* from endothelial cells, and *DEFB1* both showed a positive interaction with adipocytes, and a negative interaction effect with endothelial cells.

**Table 1:** TwinsUK macrophage proportion in adipose tissue is associated to obesity-related traits but not age. Body mass index (BMI)

| Trait | $r^2$ | $P$-value |
| --- | --- | --- |
| BMI | 0.22 | $2.2 \times 10^{-8}$ |
| Visceral Fat | 0.29 | $4.9 \times 10^{-15}$ |
| Visceral Fat (BMI adjusted) | 0.28 | $1.9 \times 10^{-9}$ |
| Android /Gynoid ratio | 0.36 | $1.2 \times 10^{-16}$ |
| Android /Gynoid ratio (BMI adjusted) | 0.35 | $1.8 \times 10^{-12}$ |
| Age | -0.02 | 0.67 |

**Table 2:** G x Cell-proportion interactions identify cell-type specific eQTLs from bulk adipose tissue gene expression profiles. First column cell-type lists the cell-type proportion estimate included in the G x Cell-proportion interaction model. Macrophage proportion interactions replicated in Fairfax et al., 2015 have proxy SNPs and stimuli condition annotated. Top three eQTL tissues in GTEx are listed based on effect size. Regulatory regions column lists HaploRegv4 annotations at the lead SNP. All promoters, enhancer and other regulatory annotation enrichments are derived from HaploRegv4.

| | SNP | Gene | $\beta$ | P-value | Fairfax et al<br>stimulus & proxy<br>SNP | GTEx top eQTL tissues | Regulatory regions |
| --- | --- | --- | --- | --- | --- | --- | --- |
| Macrophage | rs61913538 | *CLEC12A* | 0.37 | $8.4 \times 10^{-25}$ | Naïve, rs7313235 | WB, Adipose, Muscle | Blood promoter |
| | rs1063355 | *HLA-DQA1* | 0.29 | $3.0 \times 10^{-18}$ | NA | WB, Skin, Muscle | Blood promoter. Breast/skin<br>DNAase |
| | rs1351111 | *KLRC4-*<br>*KLRK1* | -0.34 | $4.6 \times 10^{-15}$ | NA | NA | Blood + skin promoter |
| | rs1351111 | *KLRK1* | -0.35 | $1.5 \times 10^{-14}$ | NA | Adipose, Fibroblast, Muscle | Blood + skin promoter |
| | rs2422631 | *SIRPB1* | -0.40 | $2.2 \times 10^{-13}$ | NA | WB, Lung, Nerve | Blood Dnase |
| | rs28383372 | *HLA-DQA2* | 0.25 | $2.2 \times 10^{-13}$ | NA | WB, Lung, Adipose | 5+ tissues promotor |
| | rs866865 | *KCNMA1* | -0.30 | $3.7 \times 10^{-13}$ | IFN-g, rs752372 | WB, Adipose, Lung | Blood enhancer |
| | rs2278589 | *MARCO* | -0.40 | $1.8 \times 10^{-12}$ | NA | Adipose, WB, skin | Adipocytes, Monocytes |
| | rs1063347 | *HLA-DQB1* | -0.29 | $7.9 \times 10^{-12}$ | NA | WB, Lung, skin | Blood promoter + histone<br>marks |
| | rs634512 | *LYZ* | -0.27 | $2.2 \times 10^{-11}$ | LPS24, rs1384 | WB, Lung, Artery | Blood promoter. Breast/skin<br>DNAase |
| | rs4528348 | *MS4A14* | 0.26 | $5.4 \times 10^{-11}$ | LPS24, rs2233253 | Adipose, Lung, Nerve | Blood promoter + enhancer<br>Liver/Lung enhancer +<br>DNAase |
| | rs2327276 | *VNN2* | -0.34 | $7.8 \times 10^{-11}$ | IFN-g, rs1883613 | Adipose, WB, Lung | Blood + skin promoter |
| Endothelial | rs28383362 | *HLA-DQA2* | -0.34 | $1.5 \times 10^{-22}$ | NA | WB, Muscle, Adipose | Blood promoter + enhancer |
| | rs1884841 | *TC2N* | -0.23 | $1.0 \times 10^{-12}$ | NA | Nerve, Adipose, Artery | Blood DNAse + enhancer |
| | rs182366 | *B3GALNT2* | 0.26 | $7.0 \times 10^{-12}$ | NA | Nerve, Adipose, Thyroid | Blood promoter, skin DNAase |
| | rs2744944 | *UHRF1BP1* | -0.26 | $8.16 \times 10^{-12}$ | NA | Artery, Muscle, Fibroblast | Blood enhancer |
| | rs2977786 | *DEFB1* | -0.27 | $4.3 \times 10^{-11}$ | NA | Adipose, Heart, Nerve | 5+ tissue promotor |
| | rs61799378 | *SLC25A24* | -0.28 | $1.5 \times 10^{-10}$ | NA | Testis, WB, Fibroblast | Blood Promoter |
| | rs4728142 | *IRF5* | -0.28 | $3.2 \times 10^{-10}$ | NA | WB, Artery, Thyroid | Blood/Fat Promoter |
| | rs9270111 | *HLA-DRB5* | 0.30 | $6.3 \times 10^{-10}$ | NA | Muscle, WB, Adipose | Blood cell enhancer |
| | rs3819715 | *HLA-DQB1* | -0.35 | $9.0 \times 10^{-10}$ | NA | Adipose, Muscle, skin | Blood cell enhancer |
| | rs3760516 | *VAMP2* | -0.33 | $9.6 \times 10^{-10}$ | NA | Nerve, Brain, thyroid | Blood promoter + enhancer |
| Adipocyte | rs28383362 | *HLA-DQA2* | 0.26 | $4.2 \times 10^{-13}$ | NA | WB, Muscle, Adipose | Blood Promoter |
| | rs2977786 | *DEFB1* | 0.27 | $4.6 \times 10^{-11}$ | NA | Adipose, Heart, Nerve | Blood Promoter |
| | rs1812350 | *B3GALNT2* | -0.25 | $6.1 \times 10^{-11}$ | NA | Nerve, Adipose, Thyroid | Blood Promoter |
| | rs1884841 | *TC2N* | 0.20 | $6.2 \times 10^{-10}$ | NA | Nerve, Adipose, Artery | Heart/muscle promoter |

Five macrophage-dependent eQTLs were replicated in a context-specific monocyte eQTL dataset (Fairfax *et al*. 2014). Four of the five were detected in a *IFN*-γ or LPS challenged state. Given macrophage proportion in adipose tissue increases with obesity, which in turn is associated with a low-level, chronic inflammatory state, macrophages in individuals with higher proportion of macrophages are also likely to be in a stimuli-responsive state. Overall, the lead G x Cell type proportion SNPs were enriched for overlap with HaploReg enhancer annotations in primary monocytes (*P* = 0.001) and neutrophils (*P* = 0.004), consistent with the large number of G x Cell eQTLs dependent on macrophage proportion (60%).

We intersected all significant G×Cell interactions with genome-wide significant (GWS) associations in the NHGRI GWAS catalogue. 9 out of 26 SNPs are GWAS variants or are in strong LD (r > 0.80, D’ > 0.9) with GWS loci for multiple immune and autoimmune disorders. 7/9 of these SNPs are within the MHC. The other two SNPs are rs1351111, which regulates *KLRK1* dependent on macrophage proportion and is coincident with GWAS lead SNPs for Behcets disease (r2 = 1; rs2617170) (Kirino et al., 2013), and rs4728142 which regulates *IRF5* dependent on endothelial proportion and is the lead SNP in GWAS’s of a range of auto-immune diseases including Inflammatory Bowel Disease (Lui et al., 2015), Ulcerative Colitis (Jostins et al., 2012), and Systemic lupus erythematous. (Han et al, 2009). *IRF5* is a transcription factor that has been implicated in the control of macrophage polarization and tissue remodelling in adipose tissue (Dalmas et al., 2015).

## Discussion

RNA-seq profiling of bulk tissue biopsies is widely used for biomarker discovery, genetics of gene expression studies, and differential expression analysis (Ren et al., 2012; Glastonbury et al., 2016; Civelek et al., 2017; Grundberg et al., 2012). but the cellular complexity of primary tissue biopsies is often unaccounted for. In this study we used *in-silico* methods to characterize the variability of adipose cell-type composition in two large bulk tissue transcriptomic datasets and explored the effects of adipose cellular heterogeneity on a range of transcriptomic analyses. Our results indicate that it is critical to account for cell-type composition when combining adipose transcriptome datasets, in co-expression analysis and in differential expression analysis with obesity-related traits.

Whilst the ability to detect interactions with estimated cell proportions is limited both in terms of sample size and the accuracy of cell type estimation from a complex tissue such as adipose, we have demonstrated its possible to detect cell-type proportion-dependent eQTLs in whole adipose tissues. We identified 26 macrophage, endothelial or adipocyte specific eQTLs within our bulk adipose tissue RNAseq datasets, and we note all of these had main-effect eQTLs in TwinsUK adipose tissue and in several GTEx tissues (**Table 2**). The presence of immune and endothelial specific eQTLs is expected in other tissues with resident immune cells and blood vessels, however, three of the four adipocyte dependent eQTLs have been found to be eQTLs in GTEx nerve tissue. Adipose tissue is spread throughout the body and around organs, and obtaining adipose-free biopsies of many tissues, including, nerve, thyroid and muscle, is technically difficult, as is clearly documented in the GTEx pathologist notes and histology slides that are provided for every biopsy. We conjecture that the presence of adipocyte-specific eQTLs in nerve tissue is a result of adipose contamination of the nerve biopsies. This suggests that estimates of tissue-sharing of expression or regulatory effects between adipose and some tissues are likely to be an over-estimate (Lui et al., 2017; Wheeler et al., 2016). Adipose contamination will also inflate estimates of tissue-shared effects between unrelated tissues if biopsies from both tissues contain adipose contamination, as will other broadly distributed ‘contaminant’ cell types such as immune cells and blood. It is thus important to consider the cell-type composition of biopsies prior to utilizing expression or eQTL data to interpret disease *loci*, and in particular before prioritizing a tissue or cell type for downstream experiments

We demonstrate that adipose cell composition is heritable and associated to body-fat distribution. Whilst some of this heritability may be mediated by overall BMI heritability which in turn drives changes in cell composition, it is possible that certain genotypes could predispose to or protect individuals from macrophage infiltration and thereby the consequences of inflammation in obesity. Heritable variability in adipocyte number could also underlie differential capacity for adipose tissue expansion and storage, which can drive ectopic fat deposition and subsequent susceptibility to downstream cardio-metabolic disease. The role of cellular heterogeneity in modulating human health and disease is a growing area of interest (Lappalainen & Greally, 2017), and further deconvolution of bulk RNA-seq datasets, aided by the expanding availability of RNA from purified cell populations and single-cell analysis, should contribute to our understanding of how genetics influence cell-type heterogeneity and its impact on health and disease.

## Acknowledgements

This study was supported by MRC Project Grant (MR/L01999X/1) to K.S.S. The TwinsUK study was funded by the Wellcome Trust and European Community’s Seventh Framework Programme (FP7/2007-2013). The TwinsUK study also receives support from the National Institute for Health Research (NIHR)-funded BioResource, Clinical Research Facility and Biomedical Research Centre based at Guy’s and St Thomas’ NHS Foundation Trust in partnership with King’s College London. This project was enabled through access to the MRC eMedLab Medical Bioinformatics infrastructure, supported by the Medical Research Council [grant number MR/L016311/1] We would like to thank Aaron Newman, the author of CIBERSORT, for significant and useful discussion.

## Author contributions

C.A.G and K.S.S conceived and designed the project. C.A.G. performed all analysis. C.A.G. & A.C.A. conducted QC. of the RNA Seq alignment. J.S.E.M. QC’d the genotype data. A.C.A. and J.S.E.M. contributed experimental and technical support as well as discussion. C.A.G. and K.S.S. wrote the manuscript. All authors read and approved the manuscript.

## Materials and correspondence

Correspondence and requests for materials should be addressed to Craig A. Glastonbury or Kerrin S. Small

## Online Methods

### RNA-seq alignment and gene quantification

All (Adipose tissue and purified cell) data were aligned, QC’d and quantified with the same pipeline to ensure comparability. Reads were aligned to the hg19 reference genome with STAR version 2.4.0.1 (Dobin et al., 2013). All aligned BAMs were then filtered to contain reads with a mapping quality greater than 10 and only reads that were properly paired and had two or fewer mismatches were kept. Samples were excluded if the failed to have more than 10 million reads map to known genes or if the sequence data did not correspond to actual genotype data as assessed using the ‘mbv’ mode of QTLtools (Delaneau et al., 2017). GENCODE annotation v19 gene counts were calculated using only protein coding genes without retained intron transcripts using featureCounts (Liao et al., 2013). All gene counts were transformed into Trimmed Mean of M-values (TMM) a unit shown to be well suited for an across cell-type study design and that also accounts for library size differences (Robinson et al., 2010; Gong & Szustakowski 2013). Whilst all protein-coding genes were used for cell type estimation (20,345 genes) as filtering lowly expressed genes could bias cell type estimates to highly abundant cells in a given tissue biopsy, genes with at least 0.5 TMM expressed in 90% of samples within a dataset were retained for transcription wide association (TWAS) and eQTL analysis.

### TwinsUK dataset

Sub-umbilical subcutaneous adipose tissue punch biopsies were collected from female twins from the TwinsUK cohort as described in Grundberg et al., 2012 and sequenced as described in Buil et al., 2015. RNAseq data is available in EGA under accession EGAS00001000805. QC of the TwinsUK genotypes has been described previously (Buil et al., 2015; Brown et al., 2015; Glastonbury et al., 2016) After QC 766 TwinsUK samples were available for analysis, of which 720 had available genotypes. The TwinsUK adipose samples had a median age of 60 [38-84] and median BMI of 25 [16-47].

### GTEx RNA-seq dataset

RNA-seq fastq data for all GTEx v6p Subcutaneous adipose tissue samples were downloaded from dbGap. GTEx subcutaneous adipose tissue samples were obtained from the lower leg of post-mortem donors. GTEx data was re-aligned and quantified using the same pipeline as TwinsUK to ensure comparability. In addition to this, gene expression PCA outliers were removed, with outliers being defined by use of k-means clustering (k=2) fit to the first 2 expression PCs. 326 QC’d samples were retained for analysis and are listed with their cell type proportions in Supplementary File 2.

### Purified cell type data

To create the adipose signature matrix, we used purified cell type RNA-seq obtained from the Sequence Read Archive (SRA) as raw fasta files. All datasets are listed in **Table S1**. One independent set of experiments were used to construct the adipose tissue signature matrix, and another independent set to construct *in-silico* simulated mixtures to test deconvolution accuracy. Purified cell type data was aligned and quantified using the same pipeline as bulk tissue to ensure comparability.

### Construction of CIBERSORT adipose signature matrix

RNA-seq obtained from cell types and their biological replicates were constructed into a purified cell type matrix with *n* rows (genes) and *m* columns (cell type). A class file, as described in Newman et al. (2015) was also constructed to describe the pairwise comparisons to perform between cell types to produce the signature matrix. The signature matrix contains all differentially expressed genes between each cell type at a specified FDR (q = 0.30, default). CIBERSORT has the additional benefit that each tissue/mixture is deconvolved with potentially different signature genes due to the algorithm implementing a ν–Support Vector Regression (ν -SVR) step, in which only the maximally separating support vectors are retained for the linear regression. SVR also aides in minimizing co-linearity as measured through the matrix condition number (κ), an ideal step when estimating cell types that are biologically closely related.

### Estimating cell types from bulk adipose tissue RNA-seq data

CIBERSORT was used to estimate cell type proportions from adipose tissue RNA-seq samples, both from TwinsUK and GTEx (Newman et al. 2015). For signature matrix construction in CIBERSORT, we used the default value of q=0.30 for the FDR because CIBERSORTs support vector regression step maximizes which variables best fit each adipose tissue mixture, so it is therefore better to have a lower false negative rate when detecting the initial set of signature genes. CIBERSORT also reports the condition number (κ) of the signature matrix, a measure of co-linearity and matrix stability. The signature matrix has a low kappa (κ = 3.22), suggesting a well-conditioned matrix was achieved. CIBERSORT provides a deconvolution P-value per sample, calculated from 1000 bootstrapped permutations (Newman et al. 2015). We required a deconvolution P-value < 0.01 corresponding to an FDR of 1%.

### *In-Silico* mixture simulations

Purified cell types were combined at random proportions to generate 1000 *in-silico* simulated cell mixtures, termed “the ground truth” (S). A mixture matrix (M) was generated by drawing variables (equal to the total number of cells to form a mixture with) from a random uniform distribution normalized to sum to one and multiplied by the purified cell matrix (C):

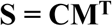

S = truth (known simulated proportions)

C = Matrix of purified cell expression profiles

M = Mixture matrix specifying amount of each cell type [0-1]

A natural amount of noise is introduced into this problem because the purified cell types are obtained from different laboratories, using different sequencing chemistries. This is ideal as the same problem is present for the deconvolution of the real Adipose tissue mixtures, making the simulated data more realistic. However, to make the problem more challenging and to assess the signature matrix’s limit and ability to deal with noise in mixture profiles, we added scaled randomly distributed Gaussian noise to each simulated sample from 10% to 100%:

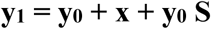

x = is a random normal variable with X ∼ N (0, 1)

y_0_ = Simulated *in-silico* mixture

y_1_ = Simulated in-silico mixture with added noise

S = scale factor [0–1]

### GTEx Histology Images and Pathologist notes

Histological images of GTEx biopsies along with accompanying pathologist notes were obtained from the GTEx web portal at https://gtexportal.org/home/histologyPage. Whilst the GTEx histology slides were prepared from a piece of material adjacent to the piece utilized for RNA-seq, they are reflective of the overall tissue sample taken.

### Heritability estimation

Heritability calculations were performed using OpenMx (Boker et al. 2011). A standard ACE model was fitted in which additive genetic, common and unique environment variance components were estimated for macrophage and adipocyte proportion between twin pairs.

### Association between cell-type composition and whole-body phenotypes

Association between cell-type proportion and whole-body phenotypes (BMI, body fat distribution and age) were conducted in the TwinsUK datasets using linear models (lm) in R. All phenotypes were collected at the time of biopsy. Body-fat distribution measurements of android, gynoid and visceral fat volume were quantified (n = 652) using dual-energy X-ray absorptiometry (DXA; Hologic QDR 4500 plus) with the standard manufacturer’s protocol.

### BMI Transcriptome Wide Association Study

Each gene expression measurement (TMM) was tested as a dependent variable in a linear mixed effects model accounted for family structure as previously described in detail (Glastonbury et al. 2016). Independent variables in addition to BMI and macrophage proportion included technical covariates that are well known to have strong effects on RNA-seq gene expression studies (Fixed effects: Insert size mode, mean GC content, Primer index) (Random effects: date of sequencing). We compared the model fit adjusted for macrophage proportion with the null model, not adjusted for macrophages, using a single degree of freedom ANOVA.

### Weighted gene co-expression network analysis (WGCNA)

Signed weighted gene co-expression network analysis was carried out using WCGNA version 1.62 (Langfelder & Horvath, 2008) in R as previously described (Langfelder & Horvath, 2008). Gene networks have been shown to follow a scale-free topology. WGCNA finds modules/clusters of highly correlated co-expressed genes using soft thresholding. The overall process has been described previously.

### *Cis*-eQTL analysis

For global *cis*-eQTL analysis, we defined each cis-window as a 1MB region around the TSS of each gene. SNPs with a MAF ≥ 5% were analysed. eigenMT was used to determine significant associations (Davis et al. 2016). EigenMT calculates the number of effective tests per *cis*-window by performing eigenvalue decomposition and taking the effective number of tests as equal to the eigenvalues that explain 99% of the variance. This procedure has been shown empirically to control the FDR similarly to permutations. All analysis was performed using inverse-rank normalized gene expression residuals corrected for experimental covariates similar to the analysis presented in Glastonbury et al., 2016. All analysis was conducted using the MatrixeQTL package (Shabalin 2012). PEER corrected residuals were obtained by correcting for 30 PEER factors (Stegle et al., 2012).

### Gene-by-environment interaction modelling

Interaction models were fitted using the ‘modellinear cross’ function in MatrixQTL (Shabalin et al., 2012). To maximize the power to detect cis-eQTLs that are dependent on cell type proportion, we inferred 30 PEER factors using inverse-rank normalized gene expression residuals corrected for sequencing date, Zygosity and family structure. Interaction models for relative macrophage proportion were adjusted for the following covariates: 30 PEER factors, mean GC content, insert size, BMI and age. Macrophage proportion was also inverse normalized to ensure normally distributed errors.

### Data Availability

TwinsUK RNAseq data is available from EGA (Accession number: EGAS00001000805). TwinsUK genotypes are available upon application to the TwinsUK cohort. GTEx data are available in dbGAP (phs000424.v7.p2). All SRA accession numbers for RNA-seq purified cell datasets for the construction of the adipose signature matrix can be found in Table S1.

