## Supplementary Materials for "Cell-type heterogeneity in adipose tissue is associated with complex traits and reveals disease-relevant cell-specific eQTLs"

***Supplementary Material***

Craig A Glastonbury[1,+†], Alexessander Couto Alves[1], Julia S. El-Sayed Moustafa[1], Kerrin S. Small[1†]

[1] Department of Twin Research and Genetic Epidemiology, King's College London, London, SE1 7EH, UK
+Current address Big Data Institute, University of Oxford, Oxford, OX3 7FZ, UK.

†Corresponding authors

| Cell type | Citation | SRA accession |
|---|---|---|
| White adipocyte | (Moisan et al. 2015) | SRR1296133, SRR1296134, SRR1296135 |
| T-cell (CD4+) | (Weinstein et al. 2014) | SRR1422906, SRR1422907, SRR1422908, SRR1422909 |
| MVEC | (DiMaio et al. 2016) | SRR2776477, SRR2776478, SRR2776479 |
| M1/M2 macrophages | (Zhang et al. 2015) | SRR2910670, SRR2910671, SRR2939145, SRR2939146, SRR2939148, SRR2939149, SRR2939150, SRR2939151, SRR2939152 |

Table S1: Purifed cell RNA-seq data from SRA used to construct Adipose tissue signature matrix

| Cell type | Adipocytes (%) | CD4+ t-cell (%) | MVEC (%) | M1 Macrophage (%) | M2 Macrophage (%) |
|---|---|---|---|---|---|
| **Adipocytes** | **99.5** | 0.003 | 0 | 0.001 | 0 |
| **CD4+ t-cell** | 0 | **100** | 0 | 0 | 0 |
| **MVEC** | 0.9 | 0 | **98.8** | 0 | 0 |
| **Macrophage** | 0 | **5.6** | 0 | **1** | **93.3** |

Table S2: Cell type (%) estimates when applying the adipose tissue signature matrix to four independent samples of purified cells. Top row represents cells present in adipose tissue signature matrix. Left most column represents independent set of purified cell RNA-seq profiles.

| Study | Cell type | Proportion (%) | Sample size | Sex | Age | Correlated with adiposity? |
|---|---|---|---|---|---|---|
| | CD4+ | 3-4.7 | 17 | M | 35-55 | n.s |
| Travers et al. (2015) | CD8+ | 0.5-5.7 | - | - | - | n.s |
| | Macrophages | 2.9 - 15.5 | - | - | - | P <0.05 (+) |
| Zimmerlin et al. (2010) | Endothelial cells | 15.4 ± 4.8 | 8 | F | - | not tested |
| Van Harmelen et al. (2003) | Adipocytes | 85 | 49 | M/F | 16-73 | P <0.05 (+) |

Table S3: Estimates from several studies that flow sorted subcutaneous adipose tissue to measure cell proportions and their relationship with adiposity.

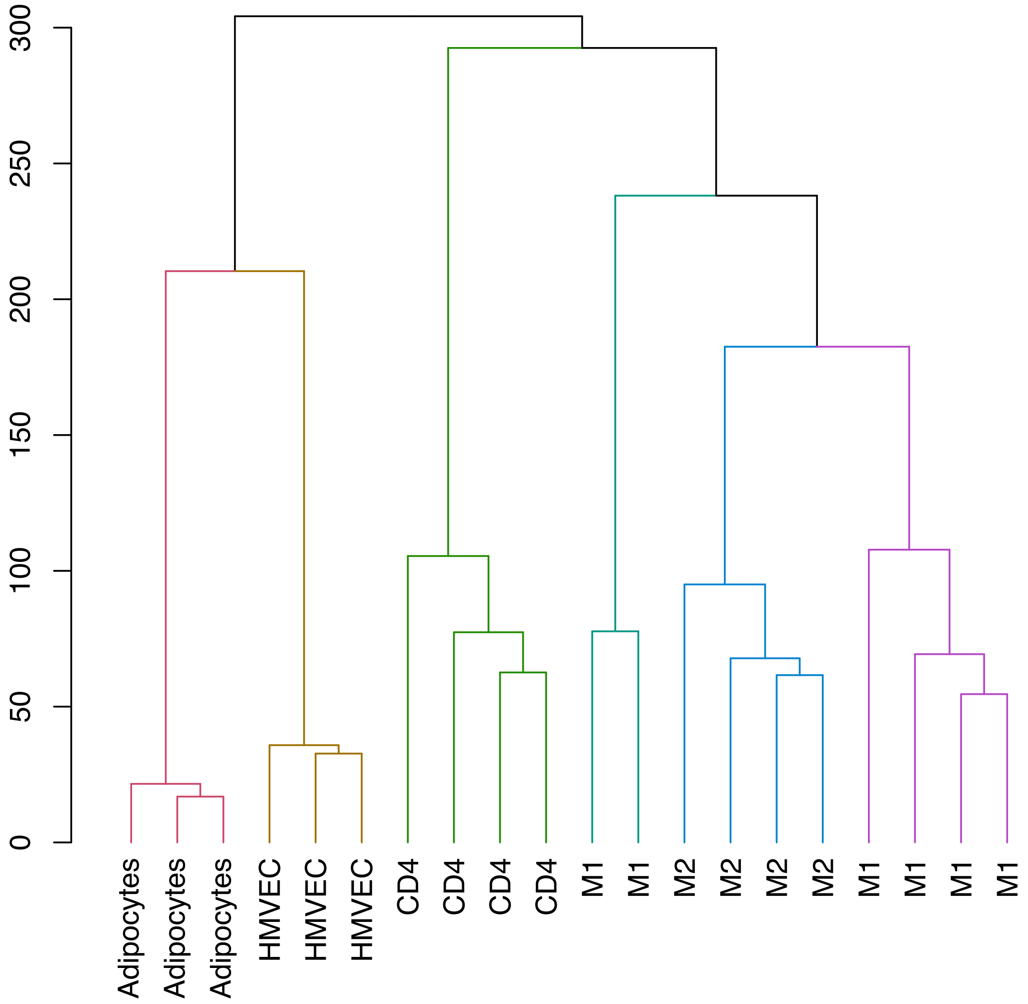


Figure S1: Hierarchical clustering of reference cells that are used to produce the signature matrix. Coloured by unsupervised k-means clustering (where k = 5). Biological hierarchy recapitulated: Non-immune (Adipocytes, MVEC) and immune cell fractions (Macrophage and CD4+ t-cells) cluster separately

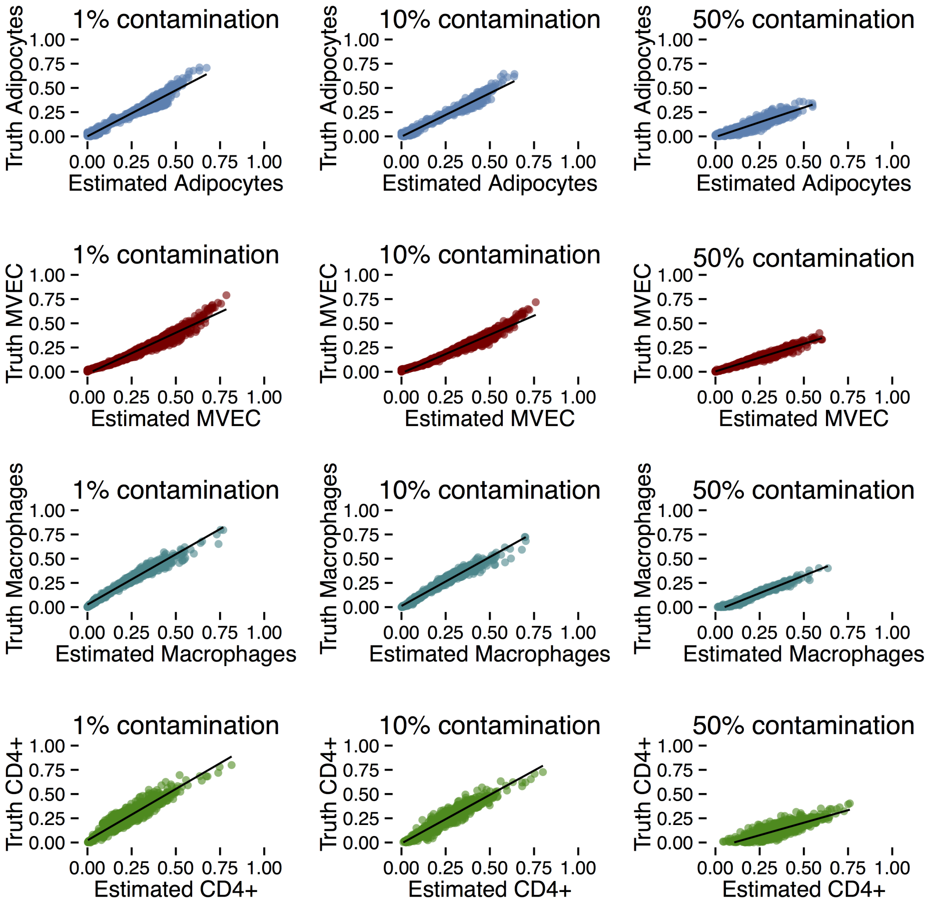


Figure S2: *In-silico* cell type estimates with unknown content added. Additional cell types known to be present in adipose tissue (Fibroblasts, Neutrophils & Dendritic Cells) but that are not estimated by the adipose signature matrix, were included in simulated adipose tissue mixtures (Adipocytes, CD4+, MVEC, Macrophages) to assess estimation accuracy with varying amounts of unknown content. The adipose tissue signature matrix is robust to unknown cell types, with cell estimates maintaining a highly linear relationship with ground truth data. An unlikely scenario of 50% unaccounted for mixture content, resulted in systematic overestimation, yet a linear relationship was still maintained.

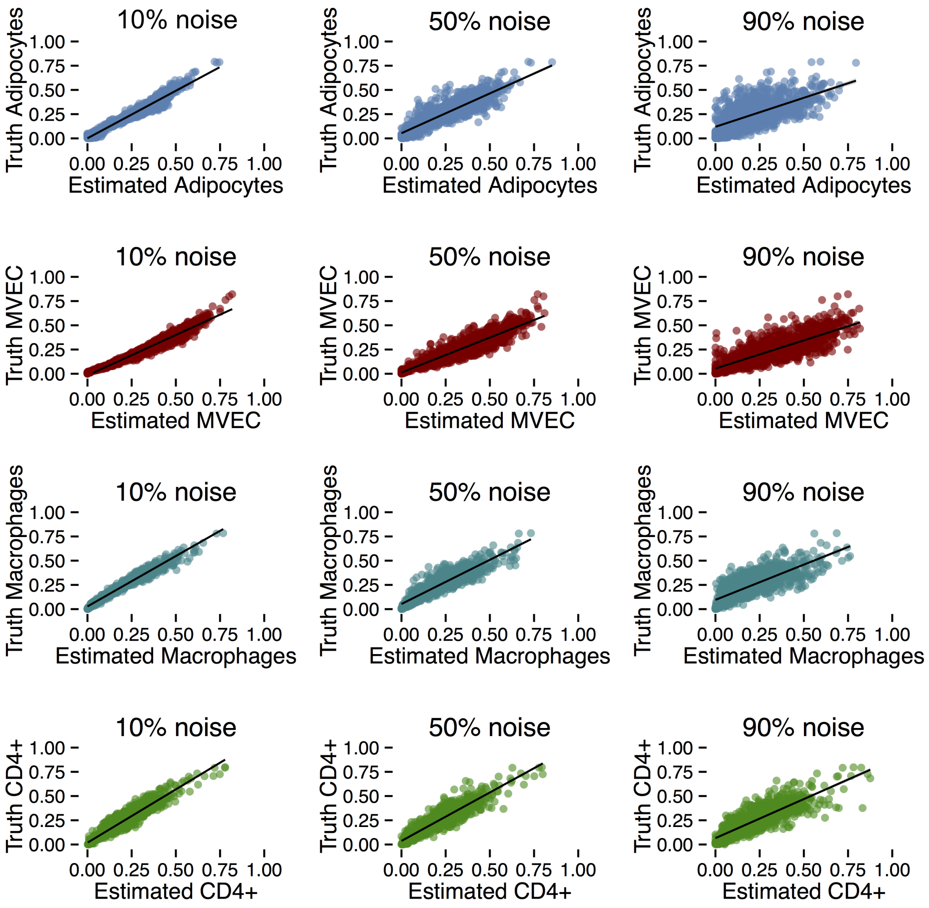


Figure S3: Cells type estimates from in-silico simulations with added scaled Gaussian noise (10, 50, 90% respectively).

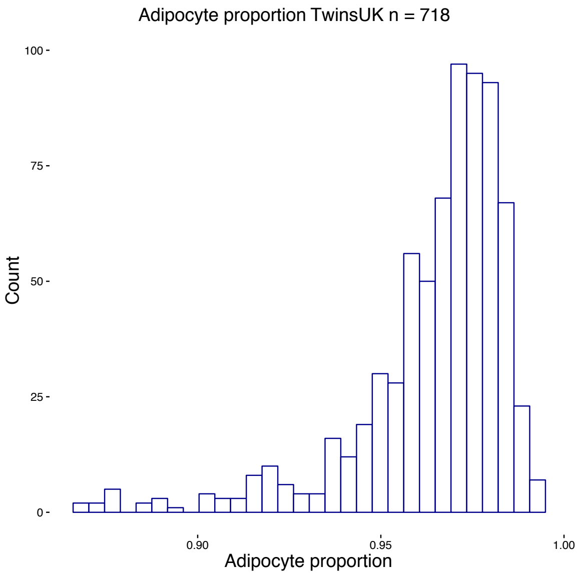


Figure S4: Estimated Adipocyte proportion in TwinsUK

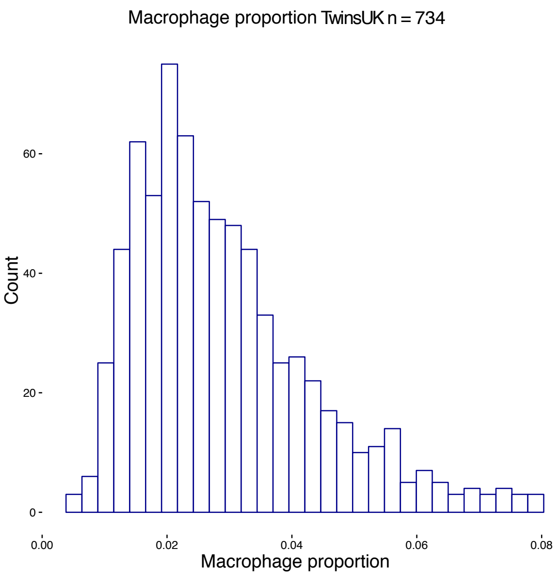


Figure S5: Estimated Macrophage proportion in TwinsUK

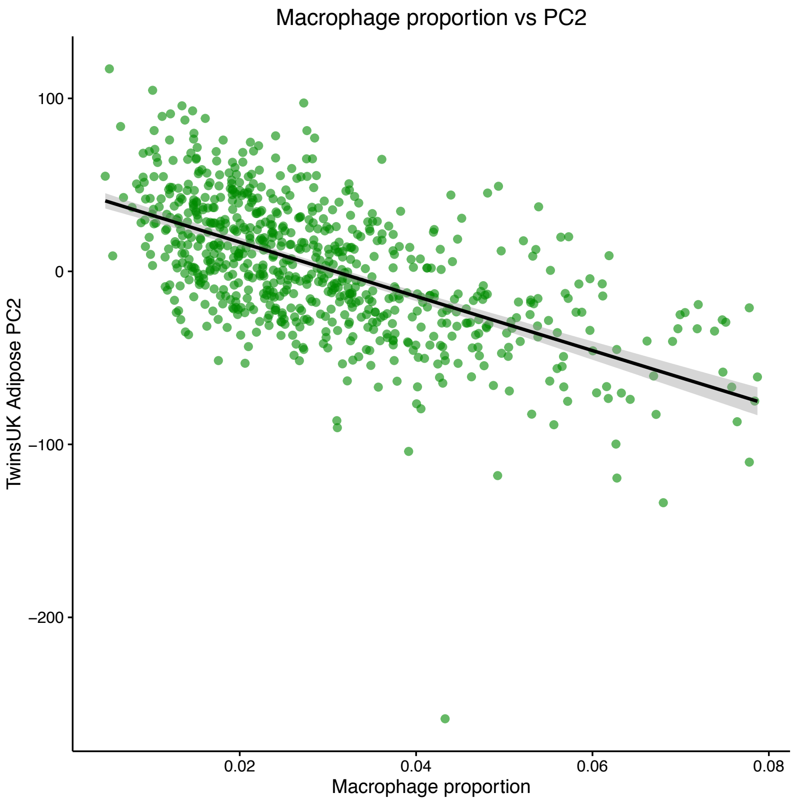


Figure S6: Adipose tissue RNA-seq PC2 captures macrophage proportion heterogenity amongst samples.

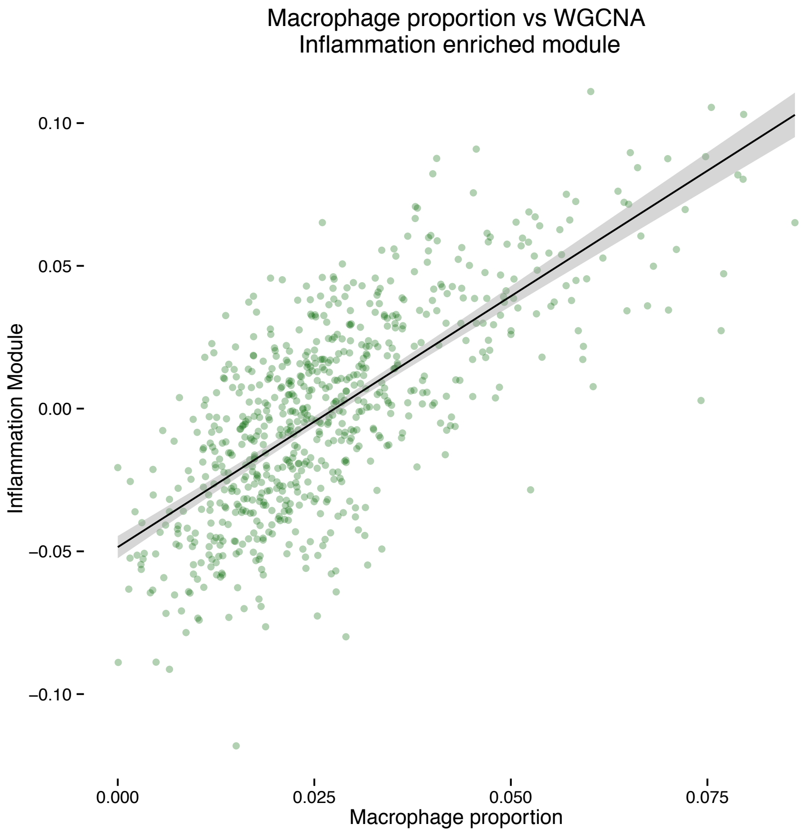


(a) Correlation of the inflammation enriched module with estimated macrophage proportion.

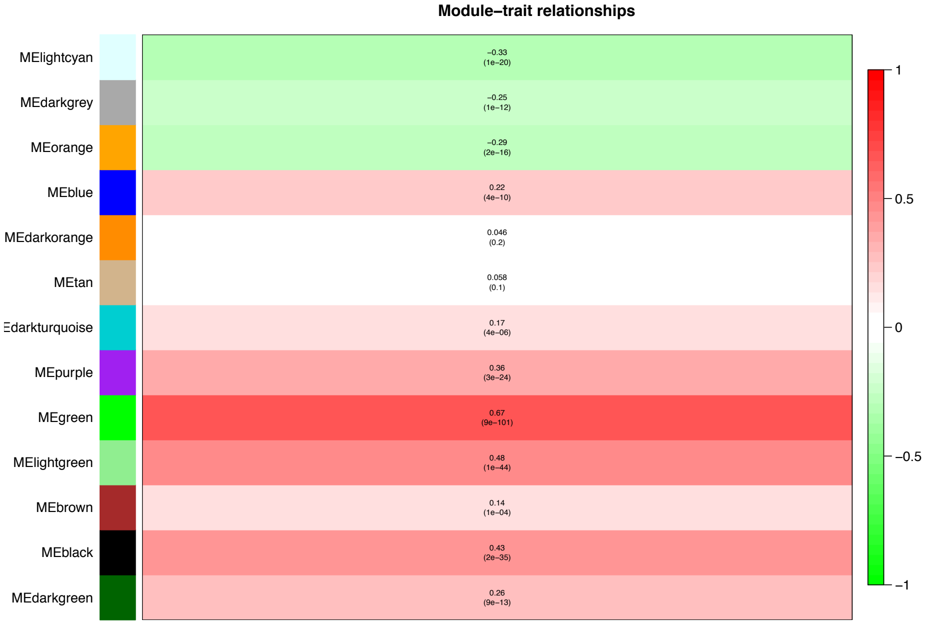


(b) Correlation of macrophage proportion with all WGCNA modules.

Figure S7: WGCNA modules and their correlation to macrophage proportion in subcutaneous adipose tissue

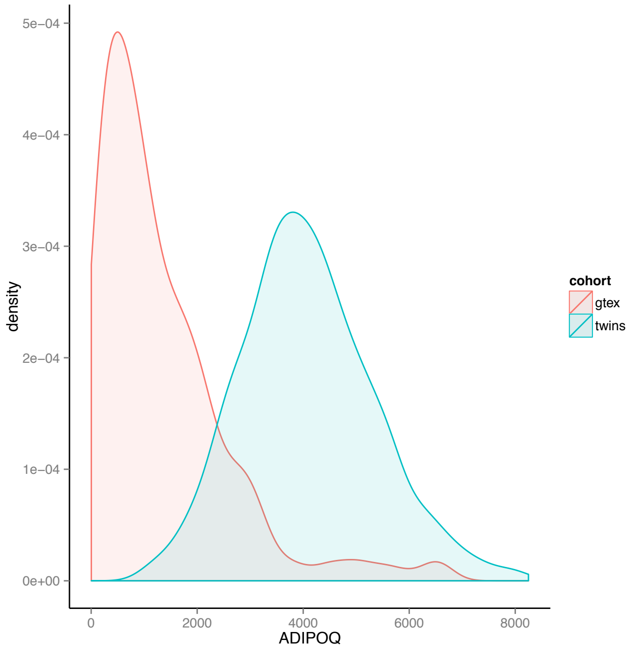


Figure S8: Distribution of ADIPOQ expression (TMM) in TwinsUK and GTEx samples.
